# An expanded view of ligandability in the allosteric enzyme PTP1B from computational reanalysis of large-scale crystallographic data

**DOI:** 10.1101/2024.01.05.574428

**Authors:** Tamar (Skaist) Mehlman, Helen M. Ginn, Daniel A. Keedy

## Abstract

The recent advent of crystallographic small-molecule fragment screening presents the opportunity to obtain unprecedented numbers of ligand-bound protein crystal structures from a single high-throughput experiment, mapping ligandability across protein surfaces and identifying useful chemical footholds for structure-based drug design. However, due to the low binding affinities of most fragments, detecting bound fragments from crystallographic datasets has been a challenge. Here we report a trove of 65 new fragment hits across 59 new liganded crystal structures for PTP1B, an “undruggable” therapeutic target enzyme for diabetes and cancer. These structures were obtained from computational analysis of data from a large crystallographic screen, demonstrating the power of this approach to elucidate many (∼50% more) “hidden” ligand-bound states of proteins. Our new structures include a fragment hit found in a novel binding site in PTP1B with a unique location relative to the active site, one that validates another new binding site recently identified by simulations, one that links adjacent allosteric sites, and, perhaps most strikingly, a fragment that induces long-range allosteric protein conformational responses via a previously unreported intramolecular conduit. Altogether, our research highlights the utility of computational analysis of crystallographic data, makes publicly available dozens of new ligand-bound structures of a high-value drug target, and identifies novel aspects of ligandability and allostery in PTP1B.

## Introduction

Rational allosteric modulation of protein function is a daunting task. To address this challenge, crystallographic small-molecule fragment screening is emerging as a powerful method to rapidly identify chemical footholds at many surface sites. An advantage of fragment screening using X-ray crystallography, as opposed to using other biophysical methods, is that it yields fully atomistic models of each fragment hit bound to the protein, which are useful for structure-based drug design of compounds that may allosterically influence distal functional sites ^1^. Crystallographic fragment screening has been used in recent years for a variety of proteins of significant biomedical interest, leading to dozens if not hundreds of hits in each case ^2–7^.

One such protein is PTP1B, the archetypal protein tyrosine phosphatase ^8^ and a highly validated therapeutic target for several human diseases including diabetes, breast cancer, and Rett syndrome ^9–13^. Previously, a large crystallographic fragment screen was performed for PTP1B: from 1966 datasets spanning 1627 unique fragments, 143 bound fragment hits across 110 structures were identified ^2^. That work leveraged an early version of the PanDDA algorithm ^14^, which exploited the density variation of ligand-soaked protein crystals, in bound and unbound states, to isolate electron density “event maps” for the ligand-bound state that enable structural modeling.

However, although it attempts to correct for global dissimilarities between structures via local real-space map alignments, PanDDA relies on a substantial degree of isomorphism between crystals to identify fragment hits ^14^. Subsequent to the original PTP1B screen ^2^, a new computational method called *cluster4x* was introduced for pre-clustering datasets using a combination of structure factors in reciprocal space and model Cα displacements in real space ^15^. A reanalysis of the original PTP1B fragment screen datasets using cluster4x revealed clustering into 17 clusters; subsequent PanDDA analyses for each cluster of datasets yielded evidence for 75 more total fragment hits, representing a substantial +52% increase in hit rate from the same data ^15^. The clustering was especially prevalent for PTP1B as opposed to other (albeit smaller) fragment screens analyzed in that study. In addition, for our subsequent rescreen of a subset of the original PTP1B fragment hits at room temperature (RT) instead of cryogenic (cryo) temperature ^2,16–20^, pre-clustering with cluster4x yielded 5 more hits (+56% increase) ^3^, further emphasizing the importance of pre-clustering fragment screen data. Unfortunately, although the overall statistics of putative additional hits from the cluster4x analysis of the original large screen of PTP1B were reported ^15^, new structural models and maps were not refined, published, or further interrogated — leaving untapped a vast resource of potential insights for a biomedically important protein system.

To address this gap, here we have modeled, refined, and analyzed 59 new crystal structures of PTP1B in complex with 65 new fragment hits, representing +54% and +46% increases respectively over the original study ^2^. This trove of new structures is now for the first time available to the public in the Protein Data Bank (PDB) ^21^, where they may be of service to those interested in phosphatase drug design and/or basic biology.

Using this wealth of data, we report several useful insights into ligandability and allosteric susceptibilities in this dynamic enzyme. Despite the seemingly thorough coverage of the protein surface with the original hits ^2^, with the new hits we observe novel binding sites, including one which has since been validated by molecular dynamics simulations and crystallography ^22^. Some other new hits uniquely bridge previously reported allosteric sites that are nearby in the structure, offering potential to develop synergistic allosteric modulators. Even when they bind in the same site as previous hits, the new fragment hits are chemically distinct, and include moieties that explore distinct structural subpockets. Intriguingly, we also identify a new fragment that provokes a previously unreported allosteric response along an internal conduit, which involves a recently reported functionally linked motif ^23,24^.

## Results

### New fragment hits from pre-clustering of X-ray datasets

To obtain dozens of previously unmodeled ligand binding events for PTP1B, we reexamined the original large-scale crystallographic small-molecule fragment screen of PTP1B ^2^. Specifically, we examined the results of a computational reanalysis of those data using the new pre-clustering software cluster4x ^15^ upstream of PanDDA analysis ^14^. The pre-clustering analysis had yielded an additional 75 putative new hits (across 72 structures), ranging from 0–14 new hits per cluster ^15^.

We reexamined the PanDDA Z-maps and event maps ^14^ from the pre-clustering analysis ^15^. Most of these maps from the pre-clustered data provide strong evidence for the presence of the soaked fragment (**Fig. S1**). Guided by these maps, we confidently modeled 65 hits (across 59 structures) out of the 75 putative new hits (across 72 structures). This resulted in a +46% increase in hits (+54% increase in structures with hits) from an already high total of 143 hits (across 110 datasets). Of the 65 modeled new hits, 50 were not originally flagged as PanDDA events; thus pre-clustering was key to identifying these hits.

For all the 59 new fragment hit structures, we have performed model validation, refinement, and deposition to the Protein Data Bank, where they are now publicly available to the broader community of scientists interested in phosphatase basic biology, drug discovery, and developing computational methods for modeling protein-ligand interactions.

### Overview of new fragment hits

Our new fragment hits are distributed across 13 distinct binding sites in PTP1B (**Fig. 1, Fig. 2a**). They have a similar distribution as the original cryogenic-temperature hits ^2^ (**Fig. 1, Fig. 2b**) and the subsequent hits from two room-temperature screens ^3^ (**Fig. S2, Fig. S3**). Many of the new hits localize to the three previously allosteric “hotspots”: the BB site, L16 site, and 197 site ^2^. No new hits bind in the active site; this is consistent with a low number of hits for the active site in the original screen ^2^, and likely is related to the charged nature of the active site and general apolar nature of the fragments. Some of our new cryo hits localize to sites that were not bound at RT (albeit with smaller fragment libraries at RT ^3^); conversely, some RT hits are at sites with no new cryo hits (**Fig. S2, Fig. S3**).

**Figure 1:**
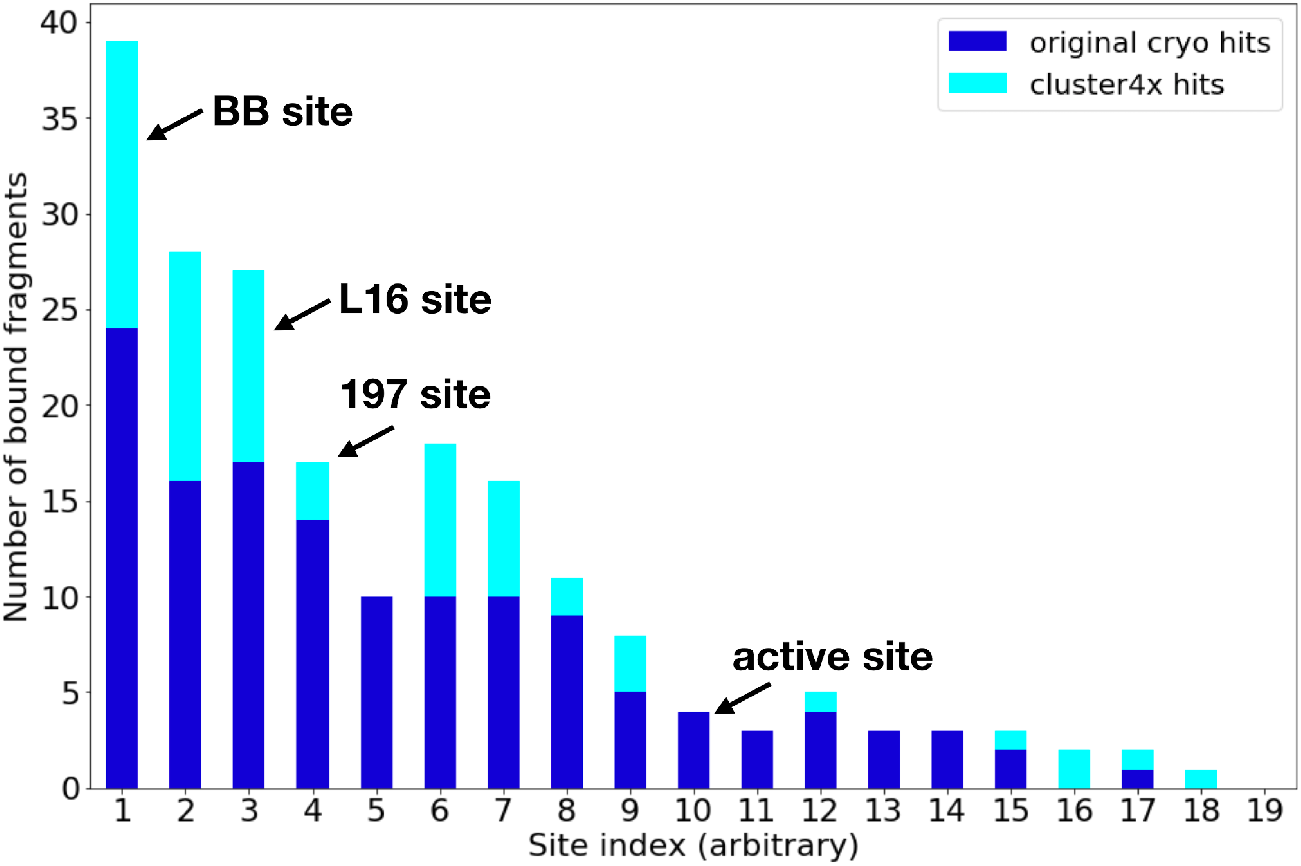
Overview of new fragment hits across binding sites. Overview of fragment hits from original cryogenic-temperature screen ^2^ and new hits reported in this study. Key sites in PTP1B, including three allosteric sites and the active site, are annotated. NB: site numbers do not coincide with previous site numbering ^2^. See also **Fig. S2**.

**Figure 2:**
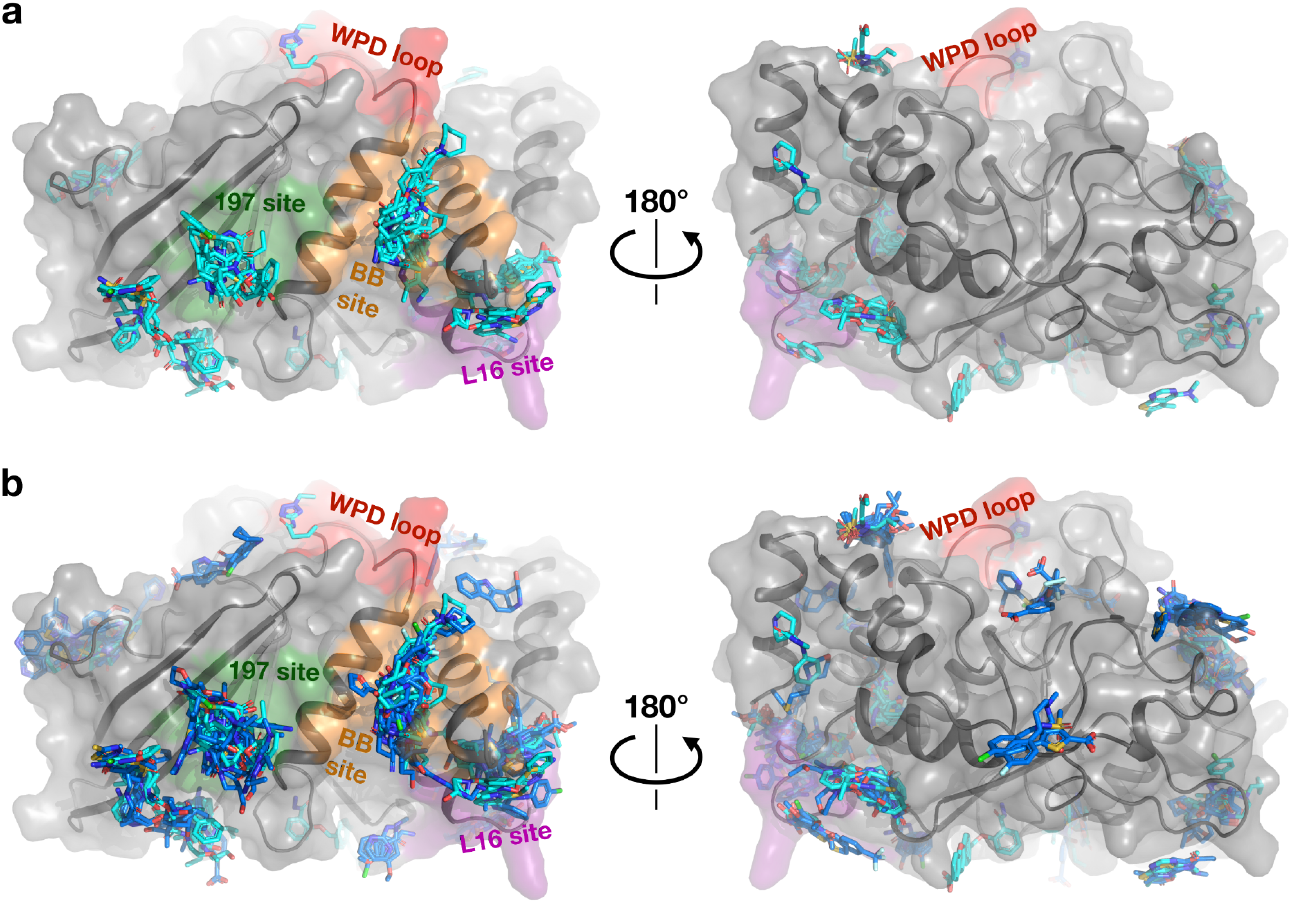
Structural overview of small-molecule fragment hits for PTP1B. a. The new fragment hits reported here (cyan) span several major binding hotspots and functional sites (green, orange, purple, and red) across the surface of PTP1B. b. Same as panel b, but with structures from the original cryogenic-temperature screen ^2^ (blue) overlaid. See also **Fig. S3**.

### New binding sites in PTP1B

The new fragment hits we report here include two new binding sites in PTP1B that were not previously seen in the original cryogenic screen or the subsequent RT screens (**Fig. 3a**). The first of these new sites is located “on top of” the catalytic WPD loop, next to the E loop (**Fig. 3b**). To our knowledge, no previous small-molecule ligands have been shown to bind to this area of PTP1B, including fortuitous “accessory ligands” such as ordered buffer components or cryoprotectants. There is a crystal contact somewhat nearby, but there is no direct crystal contact to the fragment binding site, suggesting it may also be bindable in solution. Motions of the WPD loop are critical for enzyme catalysis in PTP1B ^25^, and the WPD loop and E loop have been shown to exhibit correlated motions in the homolog HePTP ^26^, so the new binding site we report may be of interest for subsequent inhibitor development.

**Figure 3:**
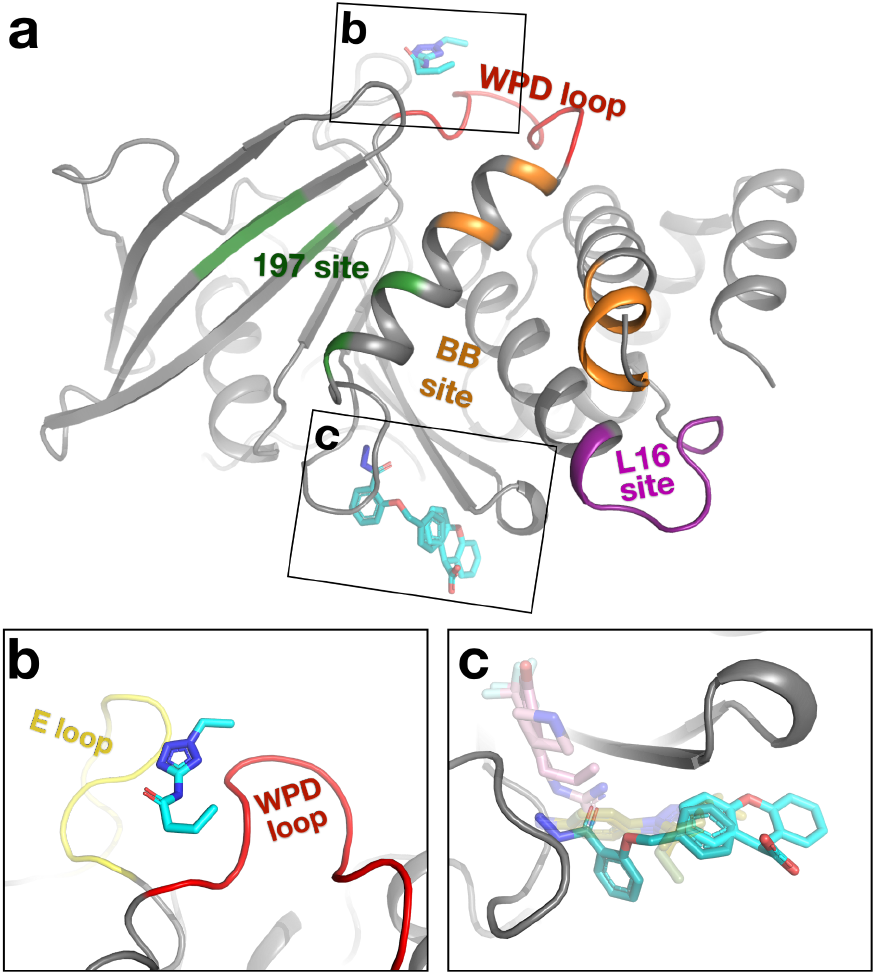
Novel binding sites in PTP1B identified from new fragment hits. Fragment hits at binding sites that were not seen in previous fragment screens of PTP1B at cryo or RT. a. Overview of binding locations. Same viewing angle and functional site coloring as **Fig. 2**, left panels. b. PDB ID 7GTQ (y0241). The fragment sits between the WPD loop (red) and E loop (yellow). c. PDB ID 7GTR, 7GTV (y0346, y1875). Crystal structures of PTP1B DES-4799 fragment series characterized by molecular dynamics simulations (PDB ID: 8g65, 8g68, 8g69) ^22^ (pale green, transparent). STEP allosteric activators (PDB ID: 6h8r, 6h8s) ^27^ (pink, transparent) are shown for context.

The second new site, with two partially overlapping new fragment hits, is located adjacent to the S loop, distal from the active site (**Fig. 3c**). Several ordered cryoprotectant molecules are also modeled in this site in other PTP1B structures, supporting its ligandability. After publication of the cluster4x reanalysis paper that first identified the new fragment hits reported here ^15^, another fragment (DES-4799) was also found to bind to this site in several poses by long-timescale molecular dynamics simulations, which, along with two analogs, was confirmed by X-ray crystallography ^22^ (**Fig. 3c**). Thus, those simulations and crystal structures provided post-facto validation of our new hits from computational reanalysis of crystallographic fragment screening data, and vice versa. Our new fragment hits and the DES-4799 series have different chemical moieties that protrude in distinct directions within the binding pocket, suggesting that medicinal chemistry efforts may fruitfully combine features of both to enhance binding at this site. Notably, these fragment poses for PTP1B also partially overlap with poses of small-molecule allosteric activators for the homolog STEP bound to the S loop region ^27^ (**Fig. 3c**), hinting at possible allosteric potential at this site in PTP1B as well.

### Increased structural and chemical diversity

The new fragment hits increase the diversity of ligand coverage of the surface of PTP1B in several distinct ways. First, some new hits bridge neighboring sites in unique ways. For example, one new hit spans two neighboring fragment-binding hotspots which are both key allosteric sites: the BB site ^2,28^ and L16 site ^2,24,29^ (**Fig. 4a**). It does so by wedging under the C-terminus of the α6 helix, which segues directly to the quasi-disordered, allosteric α7 helix ^2,30,31^. This observation raises the prospect of linking fragments, or preexisting allosteric inhibitors ^28^, in these two sites to modulate enzyme activity more potently.

**Figure 4:**
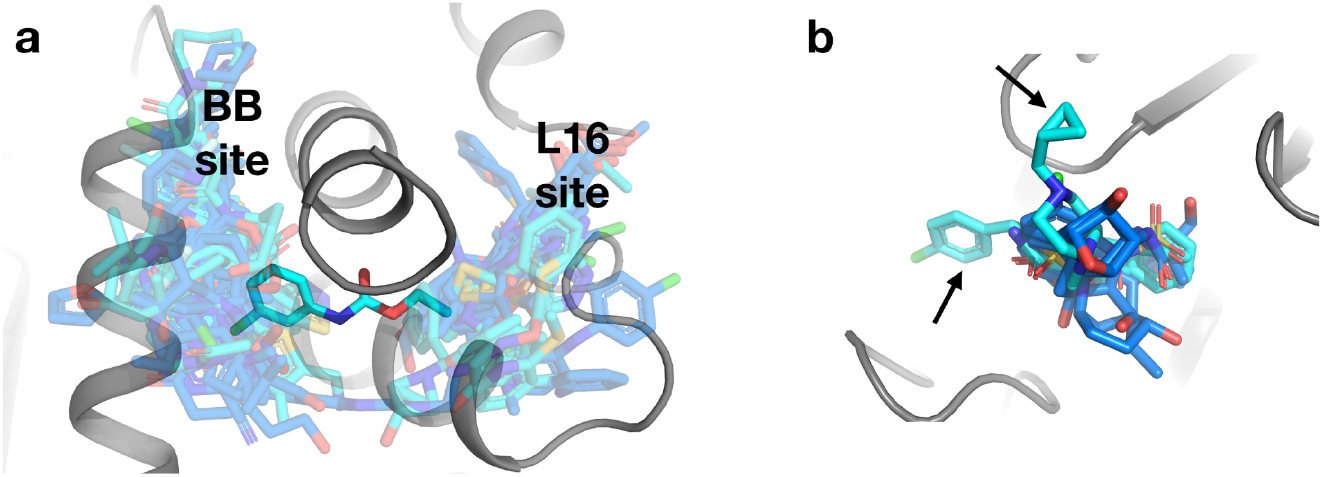
Increased coverage within and between existing binding sites. a. A new fragment hit (PDB ID 7GSA, y0288) bridges two fragment-binding hotspots in neighboring allosteric sites from the original cryo fragment hits: the BB site and the L16 site. b. In another binding site, two new fragment hits (PDB ID 7GTG, y1718; PDB ID 7GSQ, y0847) bind in poses that place chemical groups in regions of the site that were unexplored by the original cryo fragment hits (arrows).

Second, some new hits extend structural coverage within known binding sites. For example, several new hits in a site near the α2 helix protrude in different directions relative to previous hits (**Fig. 4b**).

Third, some new hits bind to new sites in addition to the original sites. Several fragments illustrate this theme, including some with new hits detected in the same structure from the same diffraction dataset, and others with new hits detected in replicate structures from different crystals soaked with the same fragment (**Fig. S4**). While in each case the fragment was already known to be able to bind to PTP1B in at least one location, these additional hits increase the chemical diversity in each binding site.

Indeed, more generally, most of the new fragment hits we report here are chemically dissimilar from other previous fragment hits within each site, as quantified by Tanimoto scores (**Fig. 5**). The low Tanimoto scores likely arise from the general dissimilarity between fragments in fragment libraries due to their small size and low complexity, in contrast to e.g. congeneric series of close derivatives of related larger compounds as might be expected further downstream in a drug development pipeline. Overall, these observations show that our new fragment hits expand the structural and chemical diversity of known ligands at many sites across the surface of PTP1B.

**Figure 5:**
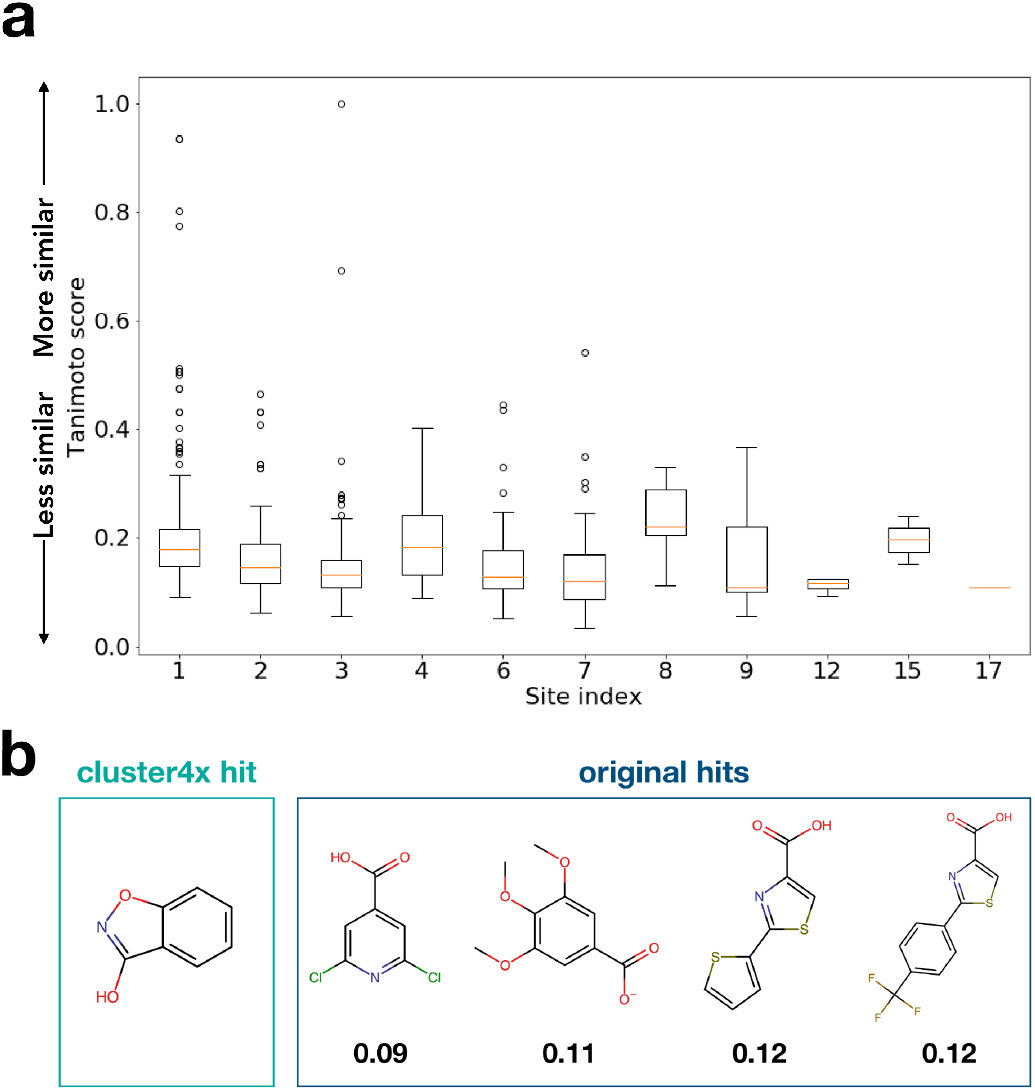
New fragment hits increase chemical diversity per binding site. a. Lowest Tanimoto score per site of any new hit to any previous cryo hit from the original screen ^2^, illustrating the extent to which the new hits represent distinct chemical scaffolds at many sites in the protein. b. Chemical structure of an example new hit (PDB ID 7GTF, y1532) compared to all the previous hits in the same site (site 12), with the Tanimoto score below each.

### Novel protein response along an allosteric conduit

In one of our new structures, we observe unanticipated conformational changes spanning 23 Å from a distal fragment binding site to the active-site WPD loop (**Fig. 6**). At the fragment site, a surface loop shifts by 0.5 Å, and a series of side chains respond in concert to the presence of the fragment. This includes the side chain of Cys226, which is immediately adjacent to Phe225 in the α4 helix, a recently discovered allosteric hub in PTP1B that we and others have recently interrogated ^23,24^; in particular, mutation of Phe225 and nearby residues surprisingly leads to increased enzyme activity ^23^, which involves the population of additional alternate conformations for several residues in this region ^24^.

**Figure 6:**
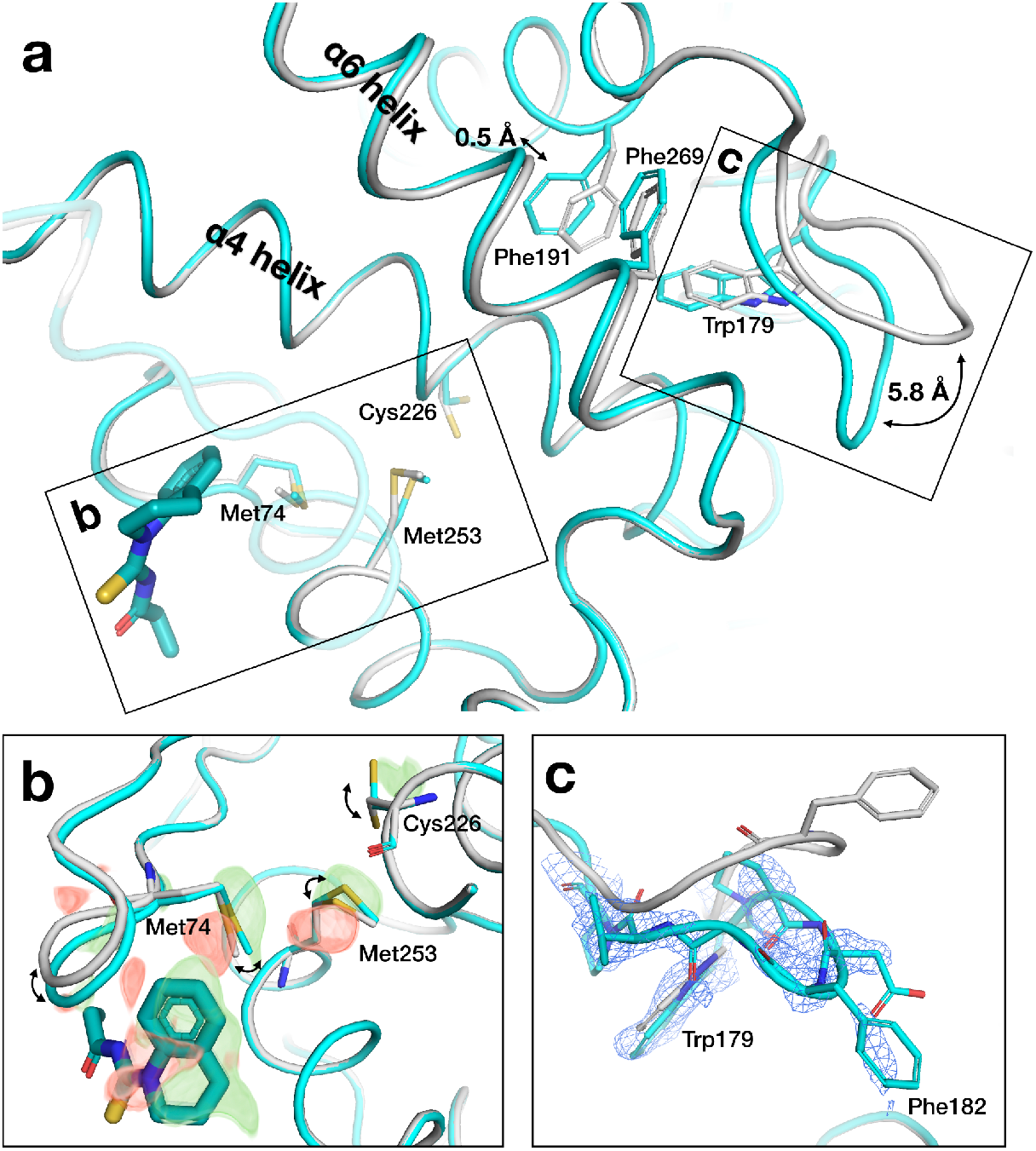
New fragment hit induces protein conformational response along allosteric conduit. a. Overview of pathway from fragment binding site to active site in PDB ID 7GTT (y0711), including ground state (gray) and bound state (cyan), with key structural elements labeled. b. Zoom-in to fragment binding site, showing conformational changes of nearby loop and series of side chains; supported by alternating difference peaks in Z-map, contoured at +/- 3 σ (green/red). c. Zoom-in to active-site WPD loop, showing shift to closed state (cyan); supported by event map, contoured at 1.5 σ.

Contacting α4, the α6 helix undergoes a concerted 0.5 Å backbone shift in response to the fragment, which is coupled to closure of the adjacent catalytic WPD loop.

There is no density evidence for oxidation of the catalytic Cys215 in the active site, arguing against the notion that chemical modification unrelated to distal fragment binding causes the WPD loop to close. Compared to the other 15 fragment hits in this site, including 10 from the original study and 5 new hits reported here, only this fragment appeared to elicit an allosteric response; the difference may be driven by the fact that this fragment protrudes the most toward the mobile loop and Met74. Taken together, we interpret these observations as allosteric response by PTP1B to a new fragment hit at a site not previously reported as allosteric, which integrates into the α-helical bundle of the tertiary structure as with other allosteric sites in this protein ^2^, but in a distinct way.

## Discussion

Crystallographic fragment screening is rapidly emerging as a powerful and accessible technique for efficiently mapping the ligandability of proteins of biomedical interest ^2–7^. Indeed, since the PanDDA method for leveraging multiple datasets to identify low-occupancy bound states was introduced in 2017 ^14^, structures from fragment screens constitute a substantial fraction (>6%) of new ligand-bound crystal structures in the Protein Data Bank. We anticipate that this proportion will only grow in the coming years, as experimental workflows at synchrotrons become more user-friendly and accessible and computational methods for bound fragment hit detection and modeling continue to mature.

One such computational advance is cluster4x ^15^, which yielded a substantial increase in fragment hits from pre-existing crystallographic fragment screen data for the therapeutic target enzyme PTP1B. Per our analysis here, cluster4x yielded 65 modelable new hits (across 59 datasets), a substantial +46% increase (+54%) over the original 143 hits (across 110 datasets). Most of these new hits bind in sites that had other fragments bound among the original hits, suggesting that, in a sense, the ligandable space of this protein had already neared saturation. However, with these new hits, we identify two new binding sites (one of which has since also been validated by another structure), one new hit that uniquely bridges neighboring allosteric sites in enticing fashion, and many new hits that explore distinct subregions of various binding sites in previously unseen ways.

In addition, with several different fragments that bind at the same non-orthosteric site, the PanDDA event maps reveal an unanticipated protein conformational response along an intramolecular pathway linking the binding site to the enzyme active site. Intriguingly, this pathway involves residues recently shown to influence catalytic function: mutations to these residues identified from coevolutionary analysis increase catalysis and decrease stability ^23,24^. These observations suggest the enticing possibility of allosteric communication between the distal binding site we highlight and the dynamic active site, ∼23 Å away. Our findings thus motivate subsequent fragment-based design ^1,32^ of higher-affinity binders at this site that may allosterically modulate function, underscoring the concept of mutations as a paradigm for drug discovery.

The 59 new ligand-bound structures of PTP1B we report here, obtained from computational reanalysis of existing experimental fragment screening data, constitute many-fold more structural information than is seen in the vast majority of publications about protein-ligand interactions. This is useful, as the rapidly emerging set of deep learning methods for protein-ligand docking and ligand design benefit from larger training sets of experimental protein-ligand structures ^33–35^; crystallographic fragment screening and improved hit detection could benefit these endeavors. On a larger scale, pharmaceutical companies house private databases that collectively contain many thousands of protein-ligand crystal structures; although making these available to developers of deep learning methods would pose a significant logistical challenge, the potential benefits to society are substantial ^34^. Other smaller-scale but practically important matters also warrant attention in this realm, including the optimal treatment of ensembles consisting of both ligand-bound and unbound states ^36^ and the need to shift away from the traditional PDB model format as the pool of unique three-letter ligand codes will soon be depleted.

Finally, all the structures we report here, and ∼94% of experimental protein-ligand crystal structures to date, are based on diffraction data collected at cryogenic temperature (<100 K). By contrast, proteins adopt distinct conformational ensembles at room temperature or physiological temperature ^2,16–20,29,37^. Importantly, so do ligands bound to proteins ^3,38,39^. The potential impact of this widespread bias in the training data for current deep learning methods for modeling protein-ligand interactions ^33–35^ or protein structures more generally ^40–42^ remains to be explored.

## Methods

### Dataset clustering and hit detection

A summary of the salient methods for the prior dataset clustering and hit detection ^15^ is as follows. Refined models and structure factors were obtained from the original cryogenic-temperature PTP1B crystallographic fragment screen ^2^ by downloading from Zenodo ^43^. Symmetry operations were adjusted per-model to ensure that all structures conformed to the same asymmetric unit. Structures exhibiting either implausibly high R values or no clear adherence to any identified cluster were excluded from the analysis. Clustering was carried out using *cluster4x* using human-guided separation on the Cα coordinates. The remaining hits formed 18 additional clusters: these consist of two sets of 9 paired clusters each, due to the Miller indexing ambiguity of (h,k,l) and (k,h,-l), which are randomly assigned during data reduction. In the previous study, all existing 110 hits separated into one pair of clusters due to the additional manual refinement of these structures which were used in the analysis. This is therefore equivalent to the common step of removing previous hits from the PanDDA run, to reduce the density variance in ligand-binding hotspots. For each cluster, Z-maps and event maps corresponding to potential ligand binding events were generated using PanDDA ^14^. This used the default parameterization for PanDDA but with a modification to reduce the minimum number of datasets to 20.

### Structural modeling and refinement

To model and refine the initial bound-state models from the initial cluster4x paper ^15^, we reexamined the PanDDA maps in detail for each example.

Fragments and associated protein changes were modeled in Coot. Some models showed structural changes like loops closing, but no ligand was seen bound in the model, so the structures were not included in this analysis. Some new binding events were found within datasets that already had ligands modeled from the original screen; in these cases, both the new and previous ligands were modeled. We avoided modeling alternate conformations for ligands to minimize model complexity.

In some cases, we modeled the WPD closed but did not model α7 ordered or L16 closed; this should not necessarily be interpreted as evidence for allosteric de-coupling, but rather is indicative of the degree of clarity at the different local areas in the event maps, which can often be noisy in places.

Waters were kept the same between the unbound and bound models, except where the PanDDA event map indicated a shift, deletion, or an addition of a new water. Ligand restraints files were calculated with eLBOW ^44^.

Because ligands are not fully occupied, to prepare for refinement we must use an ensemble of bound state plus unbound state (i.e. ground state) for refinement ^45^. We generated such an ensemble model using the giant.merge_conformations script from PanDDA 1.0.0. We then added hydrogens with Phenix ReadySet! Restraints, both between multi state occupancy groups and between local alternate locations, were generated using giant.make_restraint scripts from PanDDA 1.0.0. The argument ‘MAKE HOUT Yes’ was added to the Refmac restraint file to ensure the hydrogens were preserved.

For refinement of fragment-bound ensemble models, the published protocol for post-PanDDA refinement for deposition ^45^ was used, including the giant.quick_refine scripts from PanDDA 1.0.0 and the program Refmac ^46^. For a few examples, the script was rerun if the ligand was refined to a total occupancy greater than 1. Additionally, some hydrogens refined to 0 occupancy so they were manually edited to match the remainder of its residue. Refined bound-state models were then re-extracted using giant.split_conformations. Some examples were omitted where the ligand was not stable upon refinement.

Fragments were validated and scored using the giant.score_model script from PanDDA 1.0.0 to ensure the following five criteria were acceptable: real-space correlation coefficient (RSCC), real-space difference Z-score (RSZD), real-space observed Z-score (RSZO), ratio of ligand/protein B-factors (B-Ratio), and movement of ligand after refinement (RMSD). Outliers were removed, except some that were marginal that were visually inspected and found sufficiently supported by the density. For those with new hits in datasets that already had ligands modeled from the original screen, the original hits were kept for consistency even if their validation plots were marginal.

### Model analysis and visualization

Tanimoto scores for **Fig. 5** were calculated using RDKit ^47^ topological fingerprints, comparing the new cluster4x hits to the previous hits from the original cryo screen. PyMol was used to visualize models and electron density maps for generating figures ^48^.

## Data availability

Bound state-models, structure factors, PanDDA event maps, and traditional maps (2Fo-Fc and Fo-Fc) for all fragment-bound structures are available in the Protein Data Bank under the following PDB ID accession codes: 7GS7, 7GS8, 7GS9, 7GSA, 7GSB, 7GSC, 7GSD, 7GSE, 7GSF, 7GSG, 7GSH, 7GSI, 7GSJ, 7GSK, 7GSL, 7GSM, 7GSN, 7GSO, 7GSQ, 7GSR, 7GST, 7GSU, 7GSV, 7GSW, 7GSX, 7GSY, 7GSZ, 7GT0, 7GT1, 7GT2, 7GT3, 7GT4, 7GT5, 7GT6, 7GT7, 7GT8, 7GT9, 7GTA, 7GTB, 7GTC, 7GTD, 7GTE, 7GTF, 7GTG, 7GTH, 7GTI, 7GTJ, 7GTK, 7GTL, 7GTM, 7GTN, 7GTO, 7GTP,7GTQ, 7GTR, 7GTS, 7GTT, 7GTU, 7GTV.

For each of the 17 clusters with validated new hits, a ground-state (unbound) model is also available, along with structure factors for all datasets involved in the cluster, under the following PDB ID accession codes: 7GTW, 7GTX, 7GTY, 7GTZ, 7GU0, 7GU1, 7GU2, 7GU3, 7GU4, 7GU5, 7GU6, 7GU7, 7GU8, 7GU9, 7GUA, 7GUB, 7GUC.

In addition, we provide a Zenodo directory containing cluster identities and dataset assignments, all bound-state models, all event maps, identifying information for all fragments, and related details at https://doi.org/10.5281/zenodo.10455980.

## Acknowledgments

DAK is supported by NIH R35 GM133769.

HMG is supported by the Helmholtz Association, grant VH-NG-19-02 (Helmholtz Young Investigator Group).

We thank Jose Brandao-Neto and Justin Biel for help locating original data reduction log files and unmerged structure factor files.

