## Supplementary Figures for "An expanded view of ligandability in the allosteric enzyme PTP1B from computational reanalysis of large-scale crystallographic data"

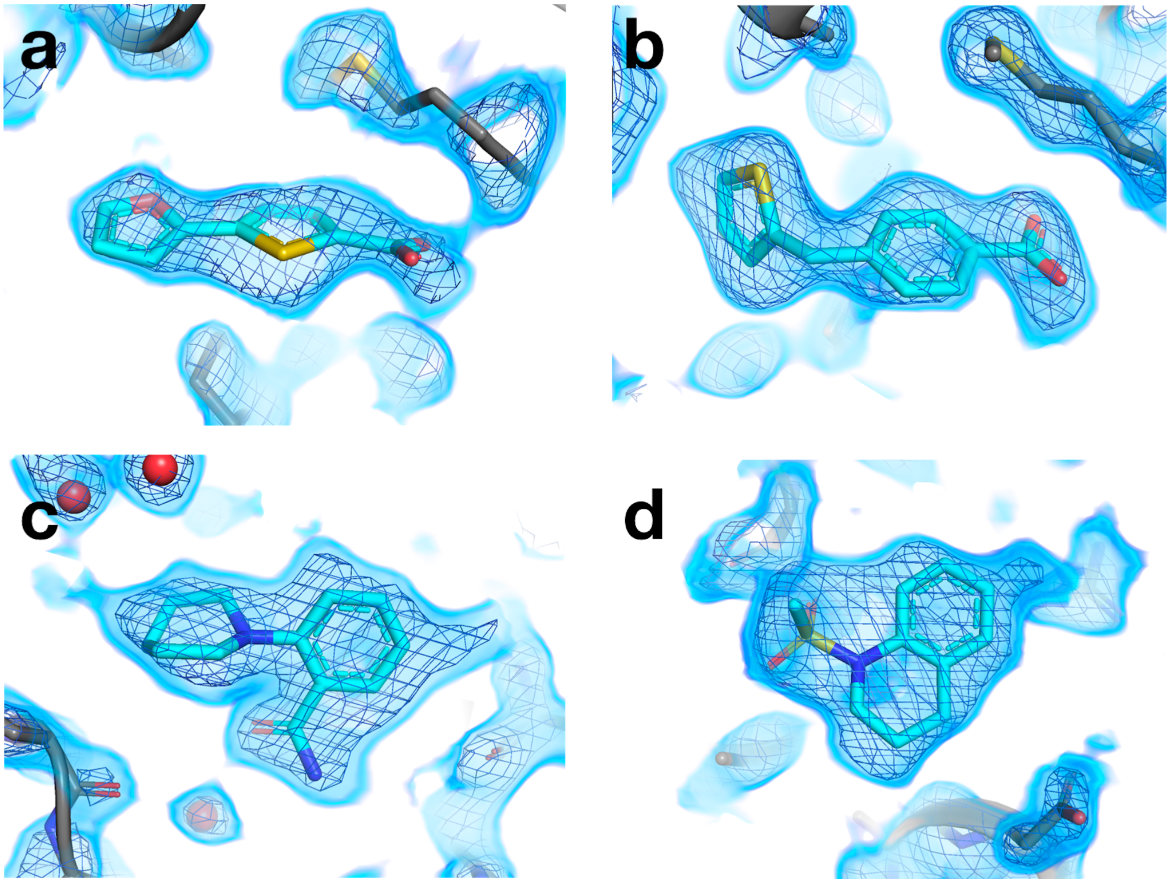


**Figure S1: Pre-clustering of datasets reveals new structures of small-molecule fragments bound to PTP1B.**

Several examples of event maps from PanDDA after cluster4x that reveal new fragment binding hits.

a. PDB ID 7GSM (y0721); event map contoured at 1.5 σ.
b. PDB ID 7GSN (y0723); event map contoured at 1.5 σ.
c. PDB ID 7GSR (y0876); event map contoured at 1.75 σ.
d. PDB ID 7GST (y0891); event map contoured at 1.75 σ.

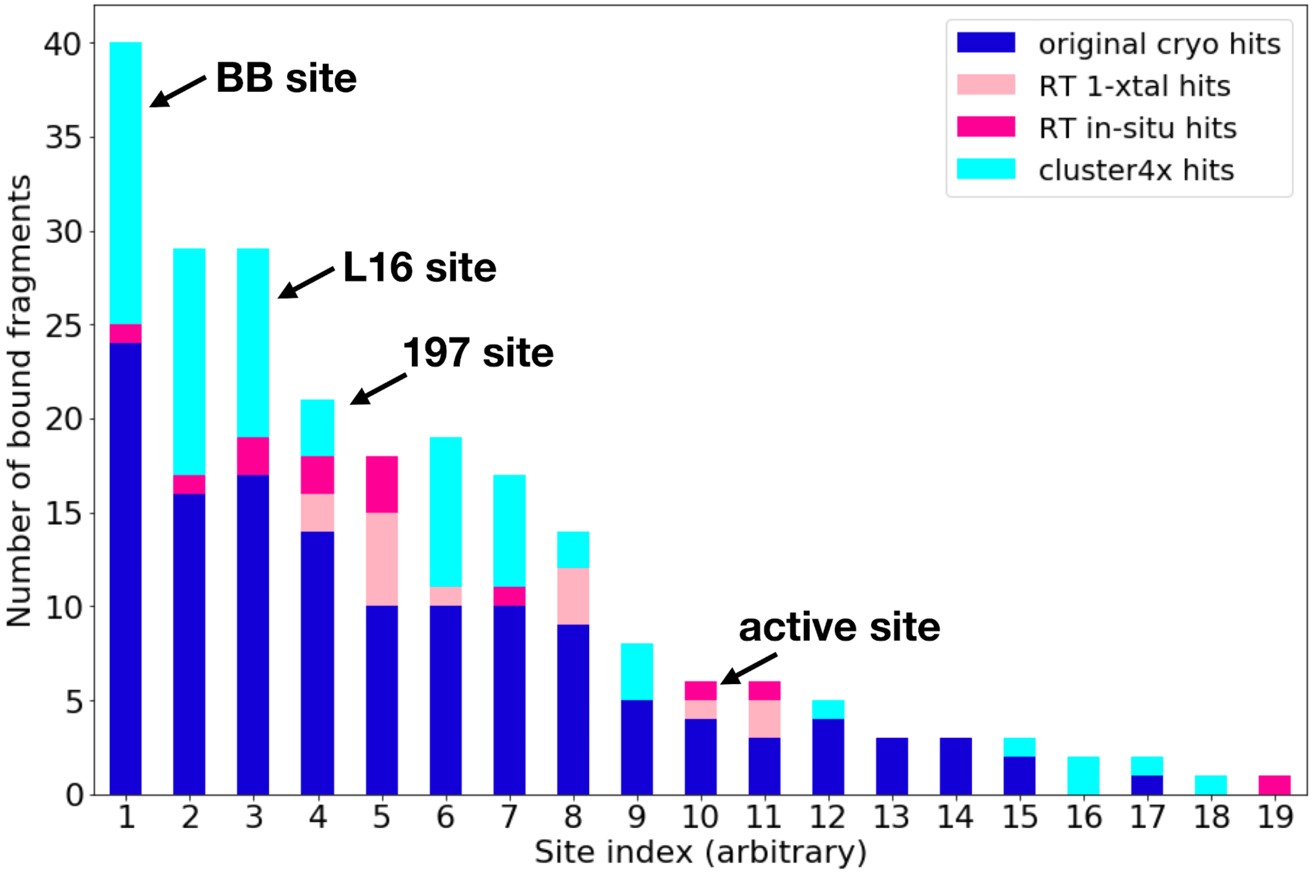


**Figure S2: Overview of new fragment hits across binding sites, compared to previous screens.** Overview of fragment hits from original cryogenic-temperature screen [2], both room-temperature (RT) screens [3], and new hits reported in this study. Key sites in PTP1B, including three allosteric sites and the active site, are annotated. NB: site numbers do not coincide with previous site numbering [2].

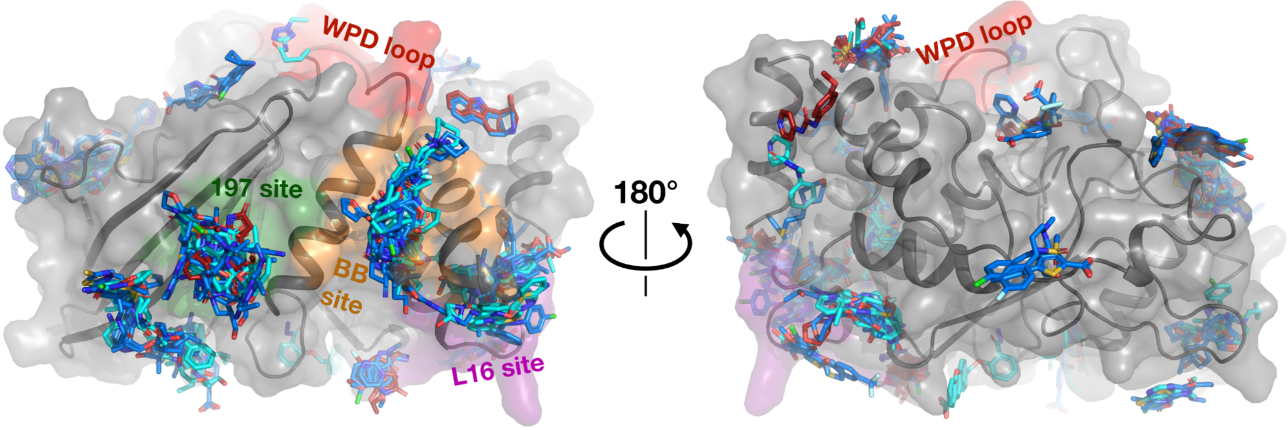


**Figure S3: Structural overview of small-molecule fragment hits for PTP1B, including room-temperature hits.**
Same as **Fig. 2**, but with structures from the original cryogenic-temperature screen [2] (blue) and room-temperature (RT) screens [3] (red) overlaid.

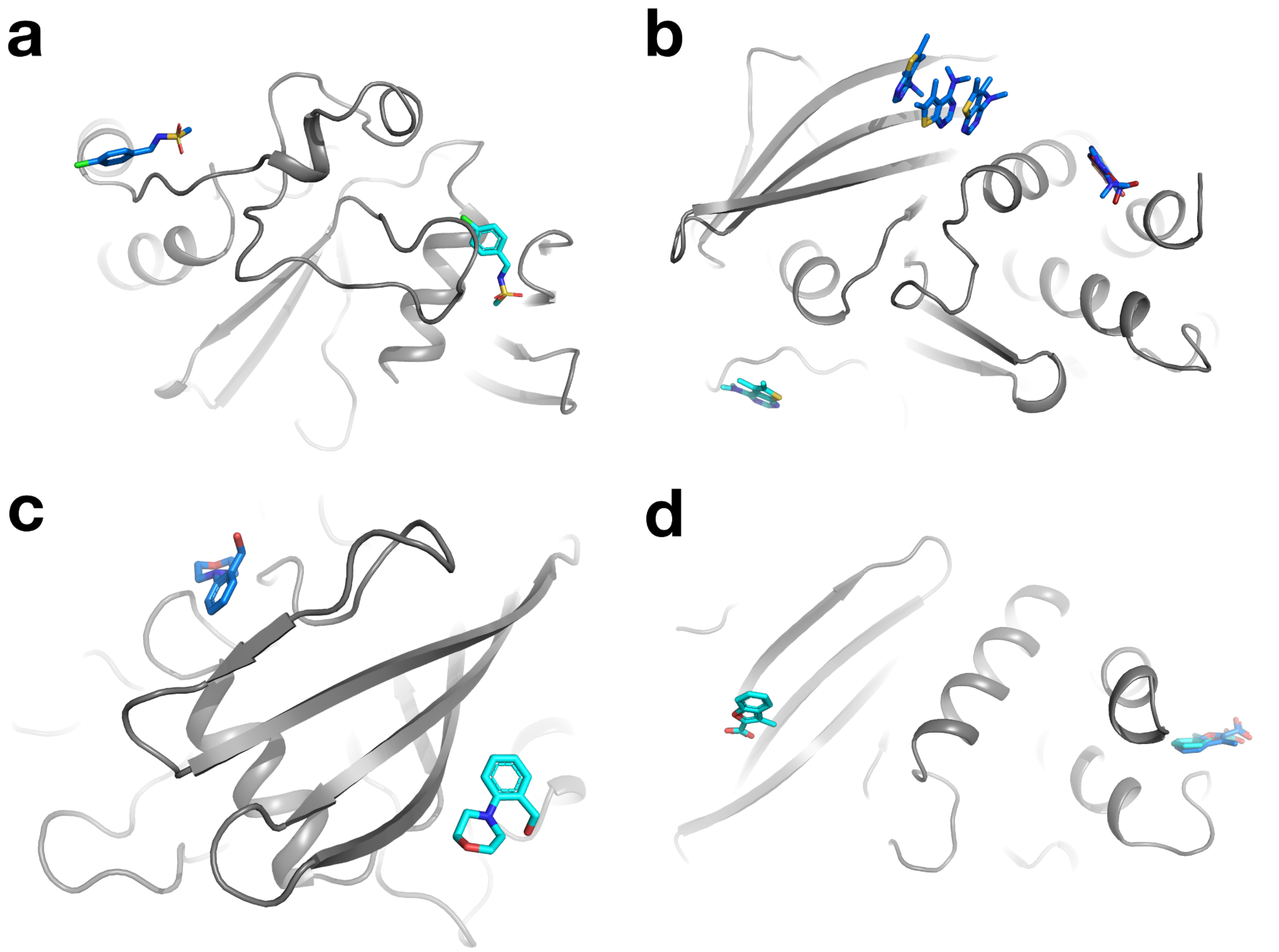


**Figure S4: Increased coverage from promiscuous fragments.**

Additional binding events for fragments with previously reported hits, now in different sites and/or datasets.

a. Same dataset, new site (PDB ID 7GSQ, y0847).
b. Same dataset, new site (PDB ID 7GS8, y0205). The new hit is distinct from the original cryo hits, which included binding at two sites: one with a single fragment molecule, and one with three fragment molecules in an artifactual stacking arrangement [2,3]. The new hit is also distinct from the single binding event for this fragment at RT [3].
c. New dataset for same fragment, new site (PDB ID 7GTJ, y1827).
d. New dataset for same fragment, same site plus new site (PDB ID 7GSX, y0986).
